# Phase-targeted Poisson closed-loop peripheral nerve stimulation modulates sensorimotor oscillations in human MEG

**DOI:** 10.64898/2026.10.01.755613

**Authors:** Stuart N. Baker, Boubker Zaaimi

## Abstract

Closed-loop brain stimulation offers a promising approach for modulating neural oscillations, but the mechanisms of phase-dependent neuromodulation remain poorly understood. We developed a real-time MEG-based closed-loop system delivering median nerve stimulation using a phase-modulated Poisson process. Sixteen healthy participants (11 female, 5 male) underwent open-loop characterization followed by closed-loop testing of eight phase-offset conditions (0°-315°), each paired with a statistically matched replay control. Closed-loop stimulation induced robust phase-dependent modulation (14.21 ± 8.24%) versus replay (7.03 ± 2.95%; p = 0.0003), with consistent phase specificity across participants (circular mean: 70.2°, referenced to each individual’s dominant frequency). At the group level, alpha oscillations (8-12 Hz) showed bidirectional modulation (mean boost: +21.0 ± 19.9% at 0°-135°; mean suppression: −13.3 ± 10.8% at 180°-315°), while beta oscillations (13-30 Hz) showed predominantly unidirectional suppression (−9.3 ± 9.6%). Successful fixed-frequency targeting required alignment with each individual’s natural peak. Misalignment led to different phase shifts producing power at different frequencies—a phenomenon we term phase-frequency coupling. Dynamic frequency tracking (8-30 Hz) automatically identified and targeted each individual’s natural peak, achieving robust bidirectional modulation without prior knowledge of peak location. These findings demonstrate that real-time phase-locked peripheral nerve stimulation selectively modulates sensorimotor oscillations beyond statistically matched open-loop stimulation. The distinct phase-excitability relationships of alpha and beta oscillations, together with the importance of aligning stimulation with individual natural peaks, establish principled guidelines for personalized closed-loop neuromodulation.

**Significance Statement:** Closed-loop brain stimulation offers a promising approach for modulating neural activity, but it remains unclear whether stimulation must be precisely timed to ongoing brain rhythms to be effective. Here, we show that real-time phase-locked peripheral nerve stimulation selectively modulates human sensorimotor oscillations, and that this effect is lost when stimulation timing is independent of brain state. By combining a novel phase-modulated Poisson stimulation paradigm with a rigorous replay control, we establish that phase alignment to the brain’s endogenous rhythm is critical for effective neuromodulation. These findings provide principled guidelines for designing personalized closed-loop neurostimulation strategies, with implications for both understanding brain state-dependent excitability and developing adaptive interventions for neurological disorders.

## INTRODUCTION

Neural oscillations play a central role in coordinating information processing across brain networks by organizing neuronal activity (Buzsáki and Draguhn, 2004; Fries, 2005). In the human sensorimotor system, 8–30 Hz activity is associated with movement preparation and somatosensory processing (Hari and Salmelin, 1997; Pfurtscheller and Lopes da Silva, 1999). The instantaneous phase of these rhythms may determine cortical excitability and synaptic plasticity, creating precise temporal windows during which neural inputs are more or less effective (Mitchell et al., 2007; Jensen and Mazaheri, 2010; Voytek et al., 2015). This principle, known as phase-dependent excitability, motivates the goal of aligning exogenous stimulation with specific phases of endogenous oscillations to modulate brain activity selectively —a paradigm known as phase-targeted closed-loop neuromodulation (Azodi-Avval and Gharabaghi, 2015; Widge et al., 2019).

However, achieving causal, phase-specific modulation presents methodological challenges. First, most closed-loop approaches rely on regular periodic stimulation, which introduces artificial rhythmicity and can entrain oscillations independently of the brain’s endogenous state, confounding interpretation (Fröhlich and McCormick, 2010; Reato et al., 2010). Pulsatile stimulation at regular intervals also has substantial power at harmonics of the fundamental frequency, which may be undesirable in some applications. Second, phase estimation typically relies on a fixed frequency band (e.g., alpha: 8–12 Hz, beta: 13–30 Hz). This fails to account for inter- and intra-individual variability in peak frequency, which can shift depending on arousal, task demands, and neural state (Klimesch, 1997; van Ede et al., 2018). As a result, stimulation may be misaligned with the individual’s true dominant rhythm, and effects attributed to ‘alpha’ or ‘beta’ phases may reflect arbitrary band definitions.

Here, we introduce a unified framework that addresses both challenges simultaneously: real-time phase-targeted stimulation using a modulated Poisson process. Instead of delivering pulses at fixed intervals, we employ Poisson-train stimulation, where inter-pulse intervals are determined randomly following an exponential distribution (Christakos, 1987; Witham et al., 2011; Koželj and Baker, 2014; Edgley et al., 2021). This stochastic pattern minimizes artificial entrainment and resembles synaptic input (Softky and Koch, 1993). Crucially, we modulate the instantaneous rate of this Poisson process in real time according to the instantaneous amplitude of phase-shifted ongoing sensorimotor oscillations. This creates an amplitude-modulated Poisson process where the probability of stimulation varies sinusoidally with oscillatory phase, allowing us to probe phase–excitability relationships without imposing rigid timing. In this study, the Poisson point process events triggered median nerve stimulation, a well-established non-invasive method for engaging somatosensory pathways and modulating cortical oscillations (Allison et al., 1989; Hamada et al., 2007; Fukuda et al., 2008; Houlgreave et al., 2022).

To implement this framework, we developed a real-time magnetoencephalography (MEG) system that dynamically tracks the dominant sensorimotor rhythm. Rather than targeting predefined alpha or beta bands, our system identifies, on each processing cycle, the frequency with the highest spectral power within the 8–30 Hz range. Stimulation is then modulated by this dynamically identified frequency. This ensures stimulation is aligned with the dominant oscillation at that moment, irrespective of whether it falls within traditional alpha, beta, or transitional ranges.

A key challenge in phase-dependent neuromodulation is distinguishing genuine phase-locked effects from arising from the statistical properties of the stimulation pattern. Even jittered or randomized open-loop stimulation can produce residual correlations with neural activity (Duchet et al., 2023). To establish causal phase-specificity, we incorporated a stringent control: each active phase-targeted condition was paired with a statistically-matched replay control. In these replay blocks, stimulation was delivered using the same phase-modulated Poisson statistics, but based on the MEG signal recorded from a previous trial, rather than the contemporary signal. This ensured identical stimulation rate distributions and temporal statistics between active and control conditions, isolating the effect of real-time phase contingency.

By integrating stochastic Poisson train stimulation, dynamic frequency tracking, and rigorous statistical controls, this study establishes a principled framework for closed-loop neuromodulation that respects the brain’s endogenous activity and provides a robust method for probing phase–excitability relationships in humans.

## Methods

### Participants

Sixteen healthy adult volunteers participated in this study (mean age 21.3 ± 2.8 years; 11 female, 5 male; 15 right-handed, 1 left-handed). All participants provided written informed consent in accordance with the Declaration of Helsinki. The study was approved by the Aston University Research Ethics Committee. Exclusion criteria included self-reported neurological disorders, implanted medical devices, or contraindications to MEG or peripheral nerve stimulation.

### Experimental Overview

The experimental design is described in detail below. The experiment consisted of two sequential phases: an open-loop characterization session to optimize stimulation parameters and derive a spatial filter, and a closed-loop intervention session to test phase-dependent neuromodulation. In the latter, participants performed a visuomotor wrist extension task while receiving closed-loop median nerve stimulation.

### Behavioural Task

One personal computer managed the behavioural task. A challenge associated with MEG recording is that all electronic equipment must be kept outside the experimental room to avoid generating noise on the MEG signal. To achieve this, we used a camera (model BFS-U3-04S2C-CS, Teledyne Flir) fitted with a telephoto lens (focal length 75 mm, Model SV-7527V, VS Technology). This was placed outside the magnetically shielded room, looking in through a side port.

A system of four mirrors was positioned within the room to allow the camera to view the hand of the participant. Mirror 1 was placed at a 45° angle in front of the side port, reflecting the camera’s view vertically upward to Mirror 2. Mirror 2, positioned higher vertically, reflected the image laterally across the room to Mirror 3, which was mounted on the side of the participant’s chair. Mirror 3 then projected the image to Mirror 4, located in front of the participant just underneath the video projector screen. Mirror 4 was angled to view the participant’s hand, which rested on a table positioned over the participant’s lap while seated in the MEG chair.

An adhesive reflective red marker (diameter 10 mm) was affixed to the tip of the middle finger of the subject’s hand, over a black glove to maximize contrast. The camera view was oriented towards the fingertips. The computer ran a customised MATLAB application, which captured frames from the camera and identified the marker position as the centre of gravity of the thresholded red camera channel. Marker position was shown as a small red circle on a video projector screen placed in front of the subject (the projector was also placed outside the shielded room, projecting in through a port).

Participants performed a visually guided wrist flexion-extension task using a cursor controlled in real time. Each trial began with the participant positioning the cursor within a bottom start target. Once the cursor was maintained within this target for 3 seconds, the middle target appeared, prompting the participant to initiate a movement toward it.

Upon entering the middle target, participants were required to hold the cursor within the target for 2 seconds, during which stimulation was delivered. Following this hold period, the target state changed, and participants were instructed to move the cursor away and subsequently return to the start position.

The task control computer was also connected to an Arduino microcontroller and digital-to-analog converter. This generated two output voltages. One signal (‘Gate’) went high during the Target 2 hold period, when stimulation should be turned on during closed loop control, and was low otherwise. The second (‘Condition’) took one of 17 different voltage levels, indicating which stimulation condition should be applied for this trial. The task computer blockwise-randomised these conditions to appear in a pseudo-random sequence. The two signals from the task computer were captured by the MEG acquisition system, making them available alongside the MEG signals for the real time control algorithm.

### MEG Data Acquisition

Neuromagnetic activity was recorded using a 306-channel Elekta Neuromag TRIUX system (102 magnetometers, 204 planar gradiometers). Data were sampled at 5000 Hz with online bandpass filtering (0.1–1650 Hz) and a 50 Hz notch filter. Participants sat upright with heads stabilized using foam padding. Head position was monitored using five head-position indicator coils placed at fiducial locations.

### Peripheral Nerve Stimulation

Right median nerve stimulation was delivered transcutaneously at the wrist using a Digitimer DS7A constant-current stimulator placed outside the shielded room. Surface electrodes (Ag/AgCl, 10 mm diameter) were placed with the cathode proximal and anode distal (∼2 cm apart). Stimulus pulses were 0.2 ms in duration. Stimulation intensity was calibrated individually by determining the motor threshold (MT), defined as the minimum current required to elicit a consistent visible thumb twitch of the abductor pollicis brevis. All subsequent stimulation intensities are expressed as percentages of this MT.

### Real-time MEG Processing System Architecture

A second personal computer ran a MATLAB script which read MEG signals in real time from the acquisition system, using the real time buffer functions of the Fieldtrip library (Oostenveld et al., 2011). Every 10 ms, this computer received an update of the last 50 sample points recorded. This included not only the MEG, but also the two voltage signals generated by the task computer. These determined whether stimulation should be given (Gate signal), and what condition should be used (Condition signal). This computer connected to a second Arduino microcontroller, which generated triggers for the median nerve stimulation.

### Open-loop Characterization Phase

Participants first underwent an open-loop session to characterize somatosensory evoked fields (SEFs) and optimize stimulation parameters. During this part of the study, participants were at rest. Stimuli were delivered as homogeneous Poisson trains at three intensities: 0%, 70%, and 100% of MT. The Poisson train was generated offline using Spike2 software (Cambridge Electronic Design) and delivered via a Micro1401 data acquisition unit. Five minutes of data were recorded for each intensity condition (around 2400 stimuli per condition).

These data were used to: (1) derive an optimal spatial filter for real-time signal extraction, (2) compute stimulus–brain coherence as a function of intensity, and (3) characterize individual sensorimotor spectral profiles to inform closed-loop targeting.

### Spatial Filter Derivation and Application

For each participant, an optimal spatial filter was derived offline from open-loop SEFs recorded at 100% MT using canonical correlation analysis (CCA). This aimed to increase the signal-to-noise ratio of the signal from sensorimotor cortex by forming an optimal linear combination of available signals. Let *X* denote a 100×204 matrix - 100 data points (15-35ms after the median nerve stimulus at 5 kHz sampling rate) x 204 signals (from the MEG planar gradiometers). Let *y* be a row vector of length 100 representing the time series from the single sensor exhibiting the maximal N20m amplitude. The CCA weight row vector *w* was obtained by maximizing the correlation:

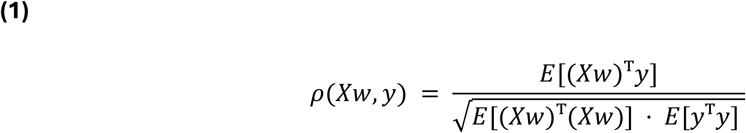

Where E(•) is the expectation operator. The derived filter weights *w* were stored and loaded during the real-time session. During closed-loop operation, the spatially filtered signal *s(t)* was obtained as:

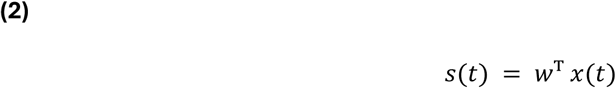

Where *x(t)* is a row vector of 204 signals at time t.

### Real-time Spectral Analysis and Peak Frequency Selection

The real-time closed-loop processing operated on a 10 ms cycle. A rolling buffer retained the most recent 0.8192 s (4096 samples) of the spatially filtered signal *s(t)*. On each cycle, the power spectral density was estimated using a Fast Fourier Transform (FFT), and the dominant frequency *f* within the 8–30 Hz range was identified.

Participants were assigned to one of three tracking conditions. For participants assigned to fixed-frequency targeting (*n* = 8), the target frequency was set to their individual alpha peak determined from open-loop recordings, with a bandwidth of ±2 Hz. For participants assigned to alpha-range targeting (*n* = 2), the target was defined as the full alpha band (8–12 Hz). For participants assigned to dynamic frequency tracking (*n* = 6), the system continuously identified the dominant frequency within the full 8–30 Hz range on each processing cycle, adapting to moment-to-moment fluctuations in the participant’s natural peak.

Rather than computing a continuous analytic phase estimate, the system employed a bank of pre-computed bandpass filters with different phase shifts and frequencies. For each target phase offset θµ and frequency *f*, a Gaussian-windowed cosine template was defined as:

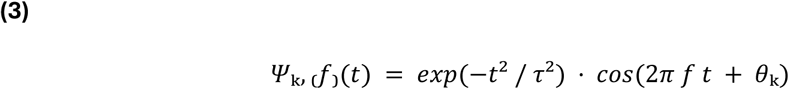

where τ = 0.8 / f. All templates were normalized to zero mean. During real-time operation, the template corresponding to the current target phase θµ and dominant frequency *f* was selected.

### Inhomogeneous Poisson Train Stimulation

The selected template was convolved with the recent signal buffer to yield a matching score *y*(*t*):

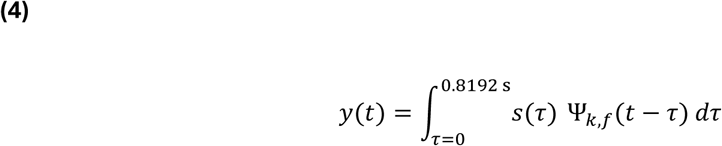

A running estimate of the standard deviation of *y(t)*, σ, was maintained over an 8 s window using a rolling buffer of recent signal values. This estimate was computed from a reference condition (with no stimulation) and applied as a global normalisation factor across all stimulation conditions. The instantaneous rate λ(t) of the Poisson process was computed as:

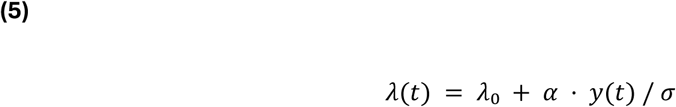

where λ。 denotes the baseline rate (10 Hz) and α the modulation gain (here set to 20 Hz).

The probability of delivering a stimulation pulse within each 10 ms processing cycle was defined as:

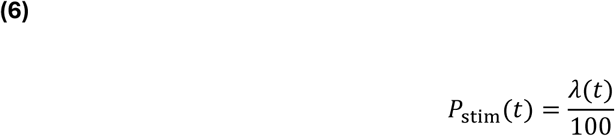

where *λ*(*t*)is the instantaneous rate from Equation (5). Within each 10 ms cycle, a pulse was delivered if a uniformly distributed random number *r* ∈ [0,1]satisfied *r* < *P*_stim_(*t*). This procedure generated a stochastic stimulation train in which stimulus probability varied with both the amplitude and phase of the MEG at the selected frequency.

For replay control blocks, instead of using the current filtered signal *s*(*t*), the signal recorded on the last trial with the same preferred phase *θ*_k_was used. This ensured identical stimulation timing statistics between active and replay conditions while removing real-time phase contingency.

### Offline Data Analysis

#### Somatosensory Evoked Fields and Source Localization

Source localization was performed on a subset of 9 participants who had complete head position data. For each participant, somatosensory evoked fields (SEFs) from the 100% MT open-loop condition were averaged over approximately 100 trials. Source reconstruction was performed using dynamic Statistical Parametric Mapping (dSPM) (Dale et al., 2000) implemented in MNE-Python. A template source space (fsaverage, oct6 spacing) and boundary element model (BEM) were used. Forward solutions were computed using individual subject head models derived from digitized head shapes. Noise covariance was estimated from the baseline period (−10 ms to 0 ms). Inverse solutions were computed with a regularization parameter λ² = 1/9. Source estimates were computed for each participant, morphed to the fsaverage template, and averaged across participants to generate a group-level source map. Anatomical labels (postcentral gyrus for S1, precentral gyrus for M1, and paracentral/superiorfrontal regions for supplementary motor area) were overlaid on the group map for visualization.

#### Stimulus–Brain Coherence

Magnitude-squared coherence between the stimulation pulse train and the spatially filtered MEG signal was computed from the open-loop characterisation data using Welch’s method. This analysis was performed at the three stimulation intensity levels (0%, 70%, and 100% of motor threshold, MT) to characterise the relationship between stimulation strength and neural entrainment at the individual participant level. These analyses were used to derive participant-specific coherence profiles which informed the selection of a common operating intensity (70% MT) for the main closed-loop experiments. Coherence was calculated for four frequency bands: theta (4-7 Hz), alpha (8-12 Hz), beta (13-30 Hz), and gamma (30-50 Hz). Coherence significance was assessed using a method based on the number of independent FFT windows (Witham and Baker, 2007). The significance limit for group-averaged coherence was calculated), with the number of windows L = 731 (5 minutes of data at 5000 Hz, 4096-point FFT, 50% overlap).

#### Analysis of Closed-loop Effects

For each participant and the 17 experimental conditions (comprising 8 closed-loop phase conditions, 8 replay phase conditions, plus one no-stimulation condition), the spatially filtered neural data from the closed-loop session were analysed. Continuous data were segmented into non-overlapping 0.8192 s epochs (4096 samples at 5000 Hz sampling rate) during the hold period of the task. The power spectrum was computed for each condition using the Fast Fourier Transform (FFT).

The frequency band of interest for power analysis was determined according to each participant’s assigned tracking method. Power within the frequency band of interest was averaged across the entire epoch for each trial.

To compute normalized power, the mean power for each stimulation condition was expressed as a percentage of the mean power during the no-stimulation baseline condition, according to:

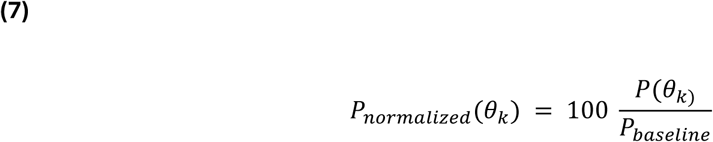

where *P(θ_k_)* is the mean power for phase condition θ_k_ (either active or replay) and *P_baseline_* is the mean power from the no-stimulation baseline condition. This yielded a percentage value for each of the eight phase bins (0°, 45°, …, 315°) for both active and replay conditions.

To quantify the amplitude of phase-dependent modulation, a Fourier-based approach was employed. For each participant and condition type (active/replay), the eight phase-power values were mean-centered by subtracting their average. The discrete Fourier transform was then computed, and the modulation amplitude was defined as:

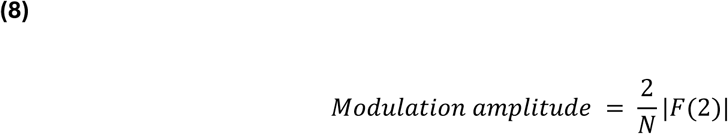

where |F(2)| is the magnitude of the second Fourier coefficient (corresponding to one sinusoidal cycle across the eight phase bins) and N = 8 is the number of phase conditions. This metric represents the peak-to-trough amplitude of the fundamental sinusoidal component of the phase-tuning curve, expressed in percentage units relative to the no-stimulation baseline.

The preferred phase for maximal power enhancement was determined from the phase angle of the second Fourier coefficient:

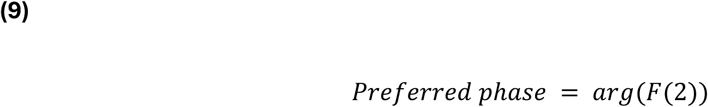

To characterize the shape of individual phase-tuning curves, cosine functions were also fitted to the eight phase-power values using nonlinear least-squares optimization, and the goodness-of-fit was assessed using the coefficient of determination (R²).

The critical comparison between active closed-loop and replay control stimulation was performed using the modulation amplitude metric. Phase-specificity at the group level was assessed by analysing the distribution of preferred phases across participants using circular statistics.

#### Phase selectivity

To quantify the concentration of phase-dependent modulation, we computed a normalized entropy-based measure. For each participant, the eight power values at different phases were normalized to sum to 1, and the Shannon entropy H was calculated. Phase selectivity was defined as 1 - H / ln(8), ranging from 0 (uniform distribution across phases) to 1 (perfect concentration at a single phase).

#### Statistical Analysis

Parametric tests were used for normally distributed data (Shapiro–Wilk test), with non-parametric alternatives applied otherwise. Effect sizes are reported as Cohen’s *d* and partial η². Multiple comparisons were controlled using false discovery rate correction (q < 0.05). Circular statistics were performed using the CircStat toolbox (Berens, 2009), with group-level phase non-uniformity assessed using Rayleigh tests.

### Code Accessibility

Custom MATLAB code used for real-time MEG processing and offline analysis is available from the corresponding author upon reasonable request.

## RESULTS

### Open-loop Characterization: Somatosensory Evoked Fields

Figure 1A shows sensor-level somatosensory evoked potentials recorded from all planar gradiometer channels for a representative participant. A clear evoked response is visible across a subset of sensors within the early post-stimulus window. The channel exhibiting the maximal response (blue trace in Fig. 1A) was used as the reference for the subsequent spatial filtering analyses.

**Figure 1.**
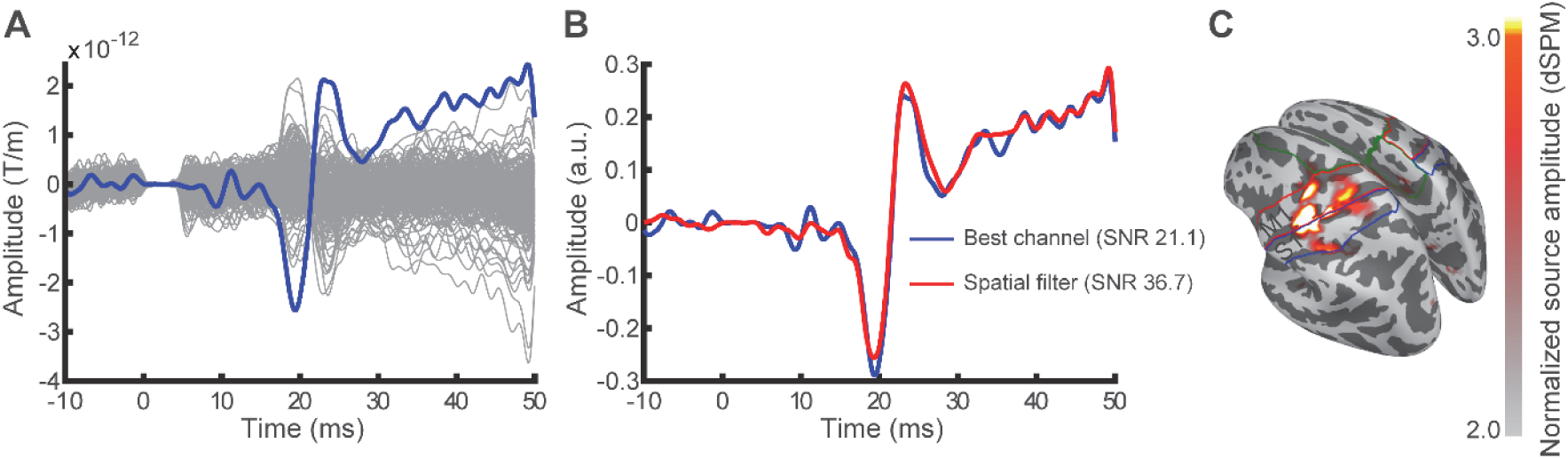
Sensor- and source-level somatosensory evoked fields (SEFs). (A) Sensor-level SEFs recorded from all planar gradiometer channels for a representative participant, with the best channel highlighted. (B) Comparison between the SEF recorded from the best individual channel (blue) and the spatially filtered SEF obtained using canonical correlation analysis (red), demonstrating an increase in signal-to-noise ratio following spatial filtering. (C) Group-level source localisation of SEFs projected onto a template cortical surface at the peak post-stimulus latency (∼20 ms), illustrating activation of both primary somatosensory cortex (S1, posterior bank) and primary motor cortex (M1, anterior bank) contralateral to the stimulated wrist. Colour bar indicates source amplitude (a.u.).

As illustrated in Figure 1B, for this representative participant the SEF recorded from the best individual channel (blue) exhibited a signal-to-noise ratio of 21.1, whereas application of the spatial filter produced a waveform with a higher signal-to-noise ratio of 36.7 (red). Across all participants (n=16), the spatial filtering procedure increased the signal-to-noise ratio from 23.1 ± 14.0 (best single channel) to 51.0 ± 24.1, an increase of 2.5 ± 1.0 times (mean ± SD).

Source localization of the somatosensory evoked field revealed activation of both the primary somatosensory cortex (S1, posterior bank of the central sulcus) and the primary motor cortex (M1, anterior bank of the central sulcus) contralateral to the stimulated wrist at the peak post-stimulus latency (∼20 ms; Figure 1C).

### Stimulus-Brain Coherence Optimization

Stimulus-brain coherence varied with stimulation intensity across frequency bands (Figure 2). Because the data were not normally distributed across all conditions (Lilliefors test, p < 0.05 for multiple comparisons), non-parametric Friedman tests were used throughout.

**Figure 2.**
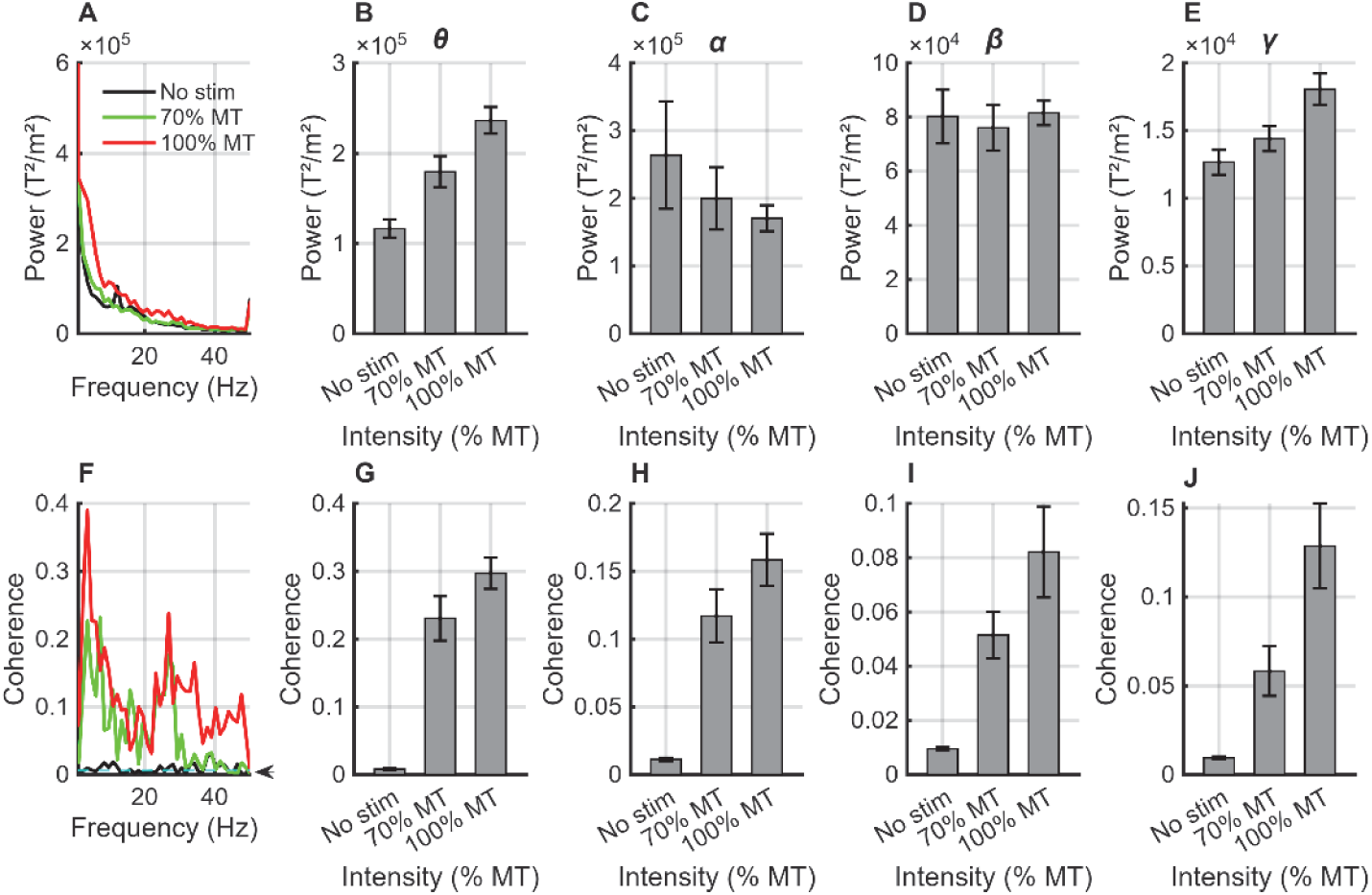
Stimulus-brain coherence dissociates from power changes. (A) Representative participant’s power spectra at 0%, 70%, and 100% MT. (B-E) Group-averaged band-specific power (n=16): theta (B), alpha (C), beta (D), gamma (E). Theta and gamma power increased significantly with intensity (Friedman test: p < 0.0001); beta power remained stable (p = 0.646). (F) Same participant’s coherence spectra with stimulus train. (G-J) Group-averaged band-specific coherence: theta (G), alpha (H), beta (I), gamma (J). All coherence bands showed significant intensity effects (Friedman tests: all p < 0.0001). The cyan dashed line and arrow in (F) indicate the significance threshold (p < 0.05). Bars represent mean ± SEM. Note the dissociation between stable beta power (D) and enhanced beta coherence (I), which increased 440.7% from 0% to 70% MT.

#### Power spectra

Theta (4-7 Hz) power increased significantly with intensity (Friedman test: χ²(2)=22.88, p < 0.0001), rising from 116,580 ± 9,969 fT²/cm² at 0% motor threshold (MT) to 179,431 ± 17,196 fT²/cm² at 70% MT (Figure 2B). Gamma (30-50 Hz) power showed similar intensity dependence (χ²(2)=22.62, p < 0.0001), increasing from 12,672 ± 918 fT²/cm² to 14,419 ± 934 fT²/cm² (Figure 2E). Alpha (8-12 Hz) power displayed a non-monotonic pattern (χ²(2)=0.12, p = 0.939), while beta (13-30 Hz) power remained stable across intensities (80,277 ± 9,949 fT²/cm² at 0% MT vs. 76,109 ± 8,519 fT²/cm² at 70% MT; χ²(2)=0.88, p = 0.646; Figure 2D).

#### Coherence spectra

Coherence spectra from a representative participant (Figure 2F) demonstrated frequency-specific peaks rather than broadband increases. Group analysis revealed significant intensity effects on coherence in all frequency bands (Figure 2G-J). Beta-band coherence (13-30 Hz) showed the most pronounced modulation, increasing from 0.010 ± 0.001 at 0% MT to 0.051 ± 0.009 at 70% MT—representing a 440.7% increase relative to baseline—and exceeded the statistical significance limit for group-averaged coherence (p < 0.05; Figure 2I). At 100% MT, beta coherence reached 0.082 ± 0.017, representing an additional 88.4% increase from the 70% level. Friedman tests confirmed significant intensity effects for all coherence bands (theta: χ²(2)=26.00, p < 0.0001; alpha: χ²(2)=23.62, p < 0.0001; beta: χ²(2)=26.00, p < 0.0001; gamma: χ²(2)=30.12, p < 0.0001). Post-hoc Wilcoxon signed-rank tests confirmed that both 70% and 100% MT produced significantly higher coherence than 0% MT (all p < 0.01). For the beta band specifically, 100% MT yielded higher coherence than 70% MT (p = 0.023).

#### Dissociation between power and coherence

The relationship between power and coherence with the stimulation train varied across frequency bands. In the theta and gamma bands, both power and coherence increased with stimulation intensity (theta power: χ²(2)=22.88, p < 0.0001; theta coherence: χ²(2)=26.00, p < 0.0001; gamma power: χ²(2)=22.62, p < 0.0001; gamma coherence: χ²(2)=30.12, p < 0.0001). In contrast, the alpha and beta bands showed a dissociation: alpha power decreased (χ²(2)=0.12, p = 0.939) while alpha coherence increased (χ²(2)=23.62, p < 0.0001); beta power remained stable (χ²(2)=0.88, p = 0.646) while beta coherence increased dramatically (χ²(2)=26.00, p < 0.0001), rising 440.7% from 0% to 70% MT. Thus, in the theta and gamma bands, stimulation amplified the existing oscillatory activity. In contrast, in the alpha and beta bands, stimulation increased the phase alignment between the stimulus train and the ongoing oscillation (coherence) without simply amplifying the oscillation itself (power). This dissociation is consistent with genuine phase entrainment rather than non-specific power amplification.

#### Individual variability and intensity selection

Individual participant analysis revealed that 75% (12/16) exhibited peak beta coherence at 100% MT, while 25% (4/16) showed optimal coherence at 70% MT. Despite the higher absolute coherence at 100% MT, we selected 70% MT for closed-loop experiments based on several considerations. First, the marginal gain from 70% to 100% MT (88.4%) was substantially smaller than from 0% to 70% MT (440.7%), suggesting diminishing returns at higher intensities. Second, 100% MT often induced discomfort that could disrupt steady-state oscillations. Third, 70% MT provided sufficient coherence (mean 0.051 over beta band) for reliable real-time phase estimation while maintaining participant comfort throughout extended recording sessions.

### Closed-loop Phase-dependent Modulation

#### Representative Single-participant Responses

Figure 3 illustrates the effects of fixed-frequency phase-targeted stimulation in a representative participant. The participant’s natural peak was in the beta range at 16.5 Hz, while the stimulation target was set to 9.8 ± 2 Hz (alpha range; Figure 3C). Time-domain traces (Figures 4A-B) show filtered neural oscillations for all eight phase conditions, with each phase (0° to 315°) represented by a distinct colour maintained across all panels. Despite the mismatch between the natural peak and the target frequency, active closed-loop stimulation (Figure 3A) produced clear phase-dependent modulation. The best boosting occurred at 90° (+116.3%), while the best suppression occurred at 315° (−3.9%), resulting in a modulation amplitude of 120.2% (Figure 3G). The sinusoidal fit (R² = 0.935) confirmed the phase-dependent nature of the modulation. In contrast, replay stimulation (Figure 3B) delivered pulses with identical timing statistics but independent of the instantaneous phase, resulting in weaker modulation (modulation amplitude: 35.3%, R² = 0.471; Figure 3H). The active/replay modulation ratio was 3.40. The phase-frequency power maps (Figures 4E-F) show normalized power as a function of both stimulation phase and frequency. Notably, the modulation effect was concentrated within the target frequency band (9.8±2 Hz), confirming spectral specificity despite the mismatch with the natural peak. The power spectra overlays (Figures 4C-D) further illustrate that active stimulation produced frequency-specific changes aligned with the target band, while replay produced weaker effects.

**Figure 3.**
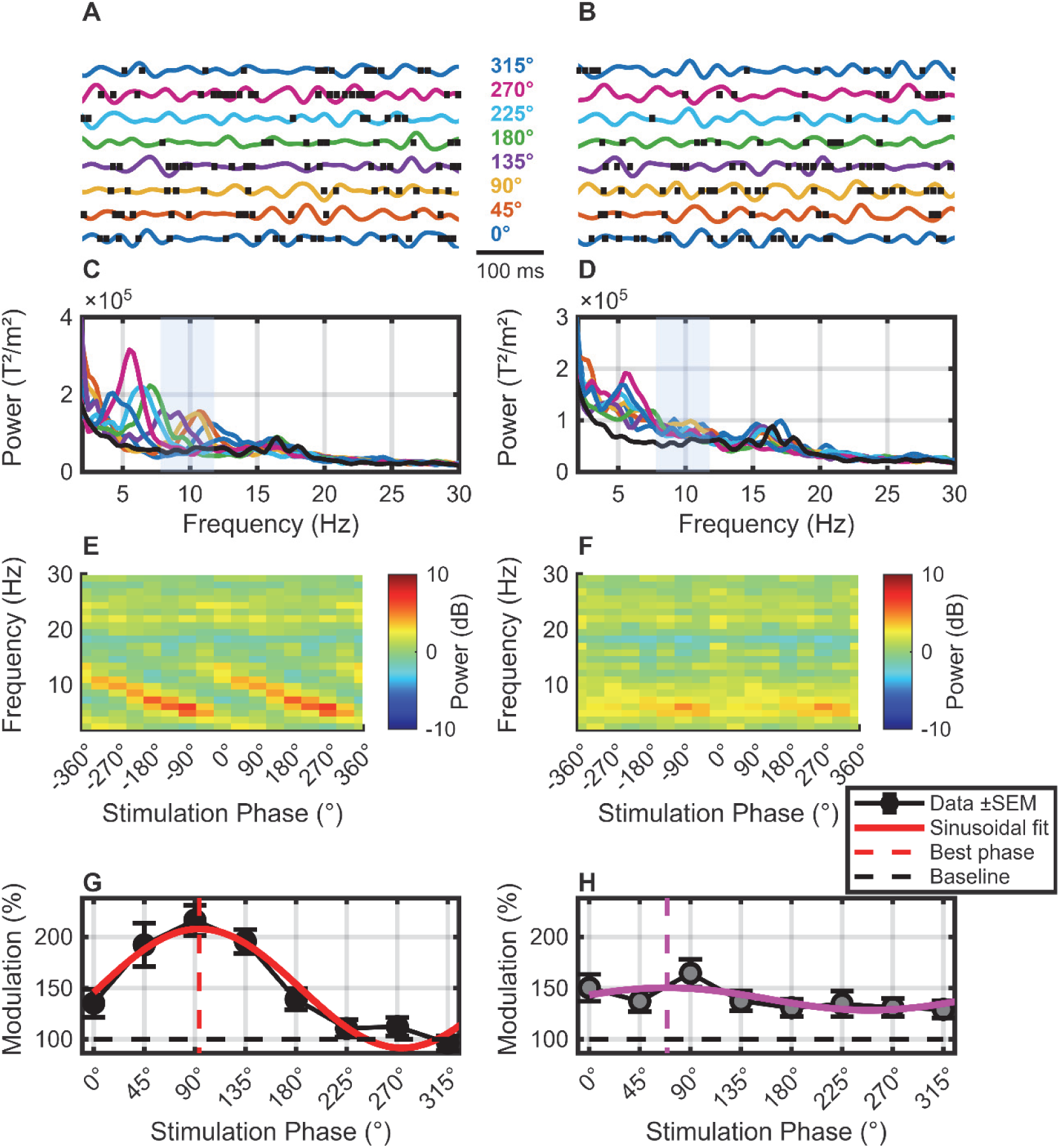
Representative single-participant analysis of fixed-frequency phase-targeted closed-loop stimulation (target: 9.8 Hz). **(A-B)** Raw time-domain neural oscillations during active (A) and replay (B) stimulation for all eight phase conditions (0° to 315°). Each phase condition is represented by a distinct colour, maintained consistently across all panels. Vertical tick marks indicate stimulation timing (active: triangles; replay: circles). Scale bar: 100 ms**. (C-D)** Power spectra for active (C) and replay (D) conditions, colour-coded by phase. The black trace indicates the no-stimulation baseline. **(E-F)** Phase-frequency power maps showing normalized power (dB relative to no-stimulation baseline) for active (E) and replay (F) conditions.**(G-H)** Phase-dependent modulation of oscillatory power within the target frequency band during active (G) and replay (H) stimulation. Points show mean normalized power (± SEM), expressed as percentage of no-stimulation baseline (100% dashed line). Red (active) and magenta (replay) curves show sinusoidal fits (active: amplitude = 120.2%, R² = 0.935; replay: amplitude = 35.3%, R² = 0.471). Red and magenta dashed lines indicate phases of maximal enhancement (active: 94.1°; replay: 69.2°). Active stimulation produced 3.40× stronger phase-dependent modulation than replay, demonstrating the efficacy of phase-locked stimulation even when the target frequency does not match the natural peak (natural peak: 16.5 Hz; C-D).

**Figure 4.**
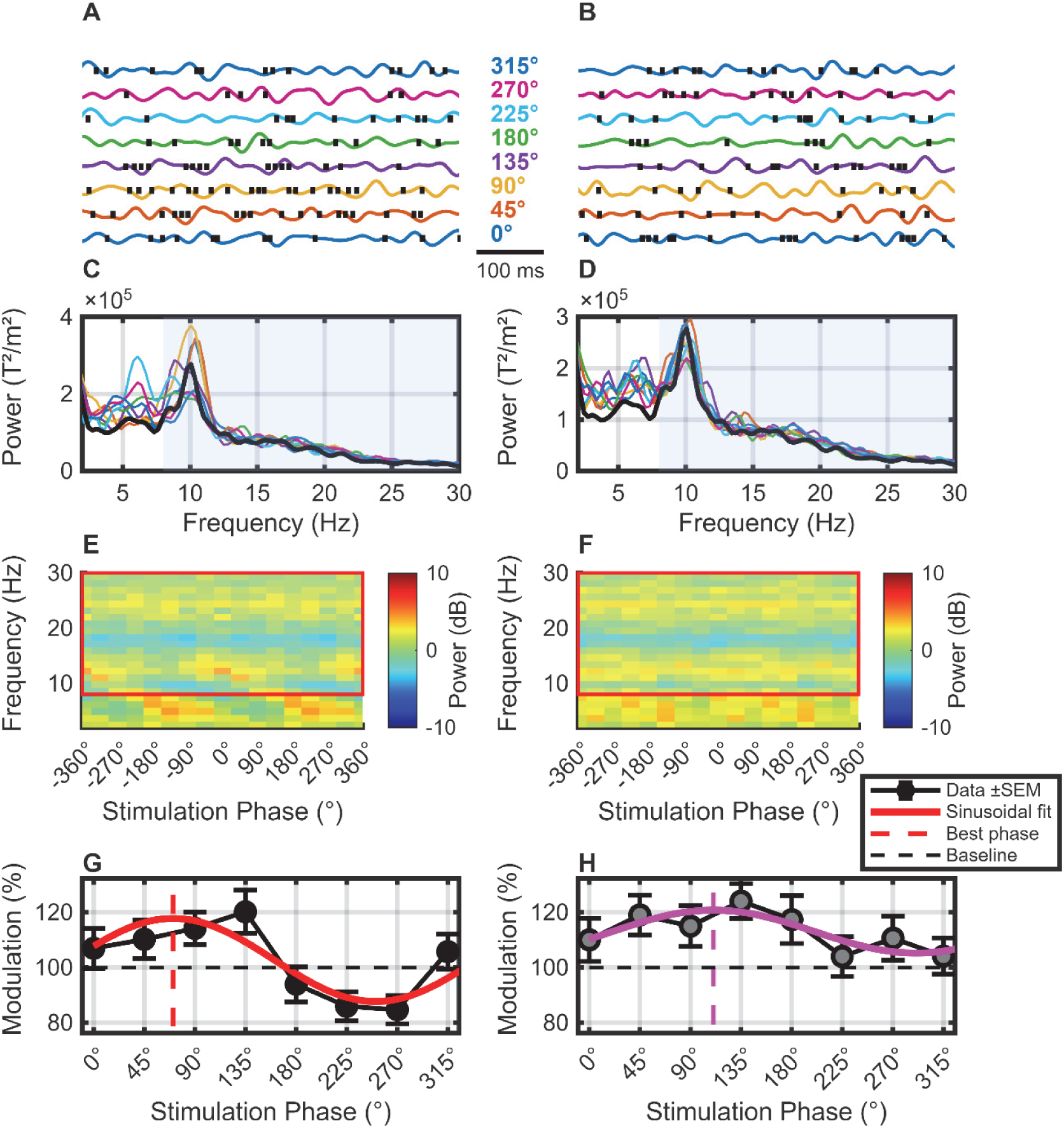
Representative single-participant analysis of dynamic frequency tracking (tracking range: 8-30 Hz). **(A-B)** Raw time-domain neural oscillations (8-30 Hz bandpass filtered) during active (A) and replay (B) stimulation for all eight phase conditions (0° to 315°). Each phase condition is represented by a distinct colour, maintained consistently across all panels. Vertical tick marks indicate stimulation timing. Scale bar: 100 ms**. (C-D)** Power spectra for active (C) and replay (D) conditions, colour-coded by phase. The black trace indicates the no-stimulation baseline. The blue shaded area highlights the dynamic tracking range (8-30 Hz) within which the system continuously estimated the dominant frequency on each processing cycle. **(E-F)** Phase-frequency power maps showing normalized power (dB relative to no-stimulation baseline) for active (E) and replay (F) conditions. Modulation was concentrated at the participant’s dominant frequency (10.1 Hz). **(G-H)** Phase-dependent modulation of oscillatory power within the 8-30 Hz band during active (G) and replay (H) stimulation. Points show mean normalized power (± SEM), expressed as percentage of no-stimulation baseline (100% dashed line). Red (active) and magenta (replay) curves show sinusoidal fits (active: amplitude = 35.5%, R² = 0.754; replay: amplitude = 20.0%, R² = 0.678). Active stimulation produced 1.77× stronger phase-dependent modulation than replay, demonstrating that dynamic frequency tracking achieves robust bidirectional modulation by continuously adapting to the dominant oscillation without requiring a priori knowledge of the natural peak location.

Figure 4 illustrates the effects of dynamic frequency tracking in a representative participant. The system continuously estimated the dominant frequency within the 8-30 Hz range on each 10 ms processing cycle and adapted the instantaneous filter accordingly (blue shaded area in Figure 4C-D). Time-domain traces (Figure 4A-B) show filtered neural oscillations for all eight phase conditions, with each phase (0° to 315°) represented by a distinct colour maintained across all panels. Active closed-loop stimulation (Figure 4A) produced clear phase-dependent modulation. The best boosting occurred at 135° (+20.2%), while the best suppression occurred at 270° (−15.3%), resulting in a modulation amplitude of 35.5% (Figure 4G). The sinusoidal fit (R² = 0.754) confirmed the phase-dependent nature of the modulation. In contrast, replay stimulation (Figure 4B) delivered pulses with identical timing statistics but independent of the instantaneous phase, resulting in weaker modulation (modulation amplitude: 20.0%, R² = 0.678; Figure 4H). The active/replay modulation ratio was 1.77. The phase-frequency power maps (Figure 4E-F) show normalized power as a function of both stimulation phase and frequency. The modulation was concentrated at the participant’s dominant frequency (10.1 Hz), demonstrating that dynamic tracking successfully locked onto the natural oscillation. The power spectra overlays (Figure 4C-D) further illustrate that active stimulation produced frequency-specific changes aligned with this peak, while replay produced weaker effects.

#### Group-level Modulation Amplitude and Phase Specificity

Figure 5 summarizes individual participant responses and the robustness of closed-loop modulation across 16 participants. Figure 5A presents representative phase-tuning curves from strong (red), medium (orange), and weak (blue) responders, defined by the top, middle, and bottom tertiles of modulation amplitude, respectively. During active closed-loop stimulation, participants exhibited clear sinusoidal phase-dependent modulation, which was reduced or absent during replay. Individual tuning curves were well described by cosine functions, with higher goodness-of-fit during active stimulation (mean R² = 0.56 ± 0.26) compared with replay (mean R² = 0.37 ± 0.20). Active stimulation produced significantly larger modulation amplitude (14.2 ± 8.2%; range: 1.5–26.7%) than replay (7.0 ± 3.0%; range: 1.5–12.1%). A paired t-test confirmed significantly stronger modulation during active versus replay stimulation (t(15) = 4.67, p = 0.0003, Cohen’s d = 1.17).

**Figure 5.**
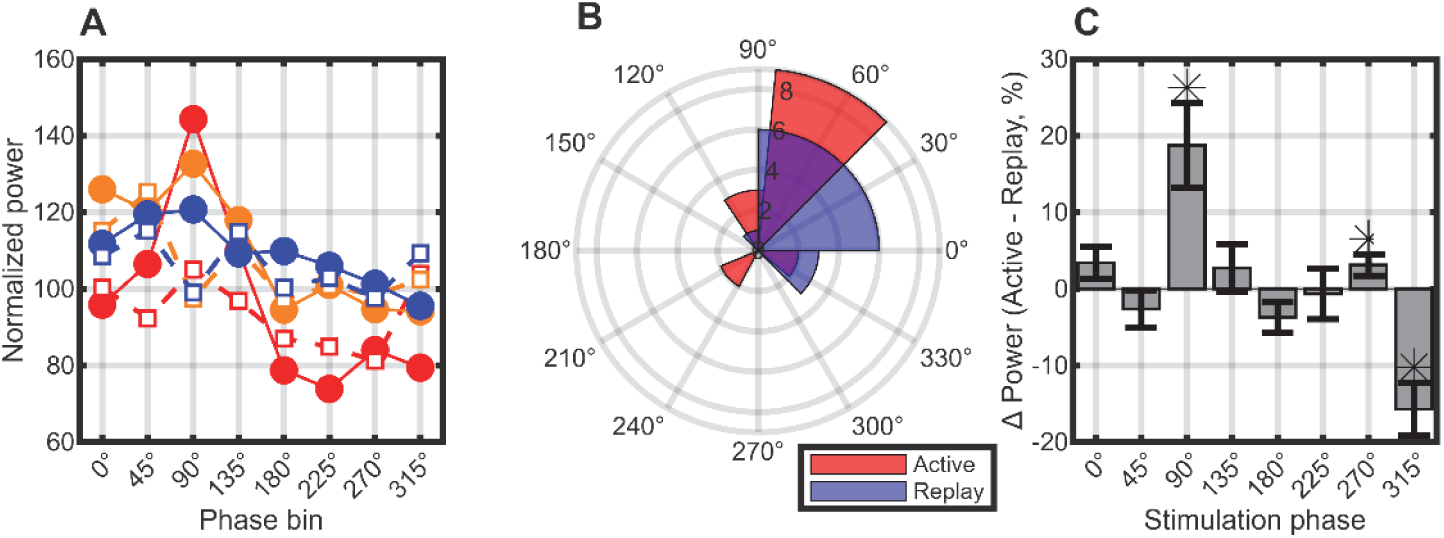
Individual tuning curves and group-level phase specificity. **(A)** Representative phase-tuning curves from strong (red), medium (orange), and weak (blue) responders, defined by the top, middle, and bottom tertiles of modulation amplitude, respectively. Solid lines indicate active stimulation; dashed lines indicate replay. **(B)** Circular distribution of the phase associated with maximal power enhancement during active (red) and replay (blue) conditions. Active and replay stimulation showed a non-uniform distribution (Rayleigh test, p = 0.0018 and p<0.0001 respectively). **(C)** Phase-resolved difference in oscillatory power (Active - Replay) within each participant’s target frequency band (see Methods). Bars show mean difference ± SEM. Asterisks indicate phases where the difference was significant (paired t-test, p < 0.05).

The phase associated with maximal power enhancement showed a non-uniform distribution across participants during active stimulation (Figure 5B; Rayleigh test, p = 0.0018, vector length r = 0.61), with a circular mean of 70.2° and circular standard deviation of 50.7°. Replay stimulation also showed a non-uniform phase distribution (p < 0.0001, r = 0.86), but with weaker modulation amplitudes, reflecting residual structure imposed by the stimulation waveform rather than genuine phase-locked interaction.

Phase-resolved differences between active and replay conditions are shown in Figure 5C. Active stimulation produced systematic phase-dependent enhancement relative to replay, with significant differences at 90° (mean difference: 18.75%, p = 0.0041), 270° (mean difference: 3.13%, p = 0.0411), and 315° (mean difference: −15.69%, p = 0.0004; paired t-test).

### Individual Tuning Curves and Phase Selectivity

Figure 6 examines the relationship between modulation strength and tuning consistency. Figure 6A shows that modulation amplitude was positively correlated with cosine fit quality for both active (r = 0.73, p = 0.001) and replay (r = 0.91, p < 0.001) conditions, indicating that larger effects were accompanied by more coherent phase tuning. Figure 6B compares individual modulation amplitudes between active and replay conditions. Most participants fell below the identity line, indicating stronger modulation during active stimulation. The correlation between active and replay modulation was r = 0.80 (p < 0.001). Figure 6C demonstrates that phase selectivity, quantified as 1 – normalized entropy, was positively correlated with modulation amplitude (r = 0.72, p = 0.002), indicating that stronger closed-loop effects were accompanied by more concentrated phase-specific modulation.

**Figure 6.**
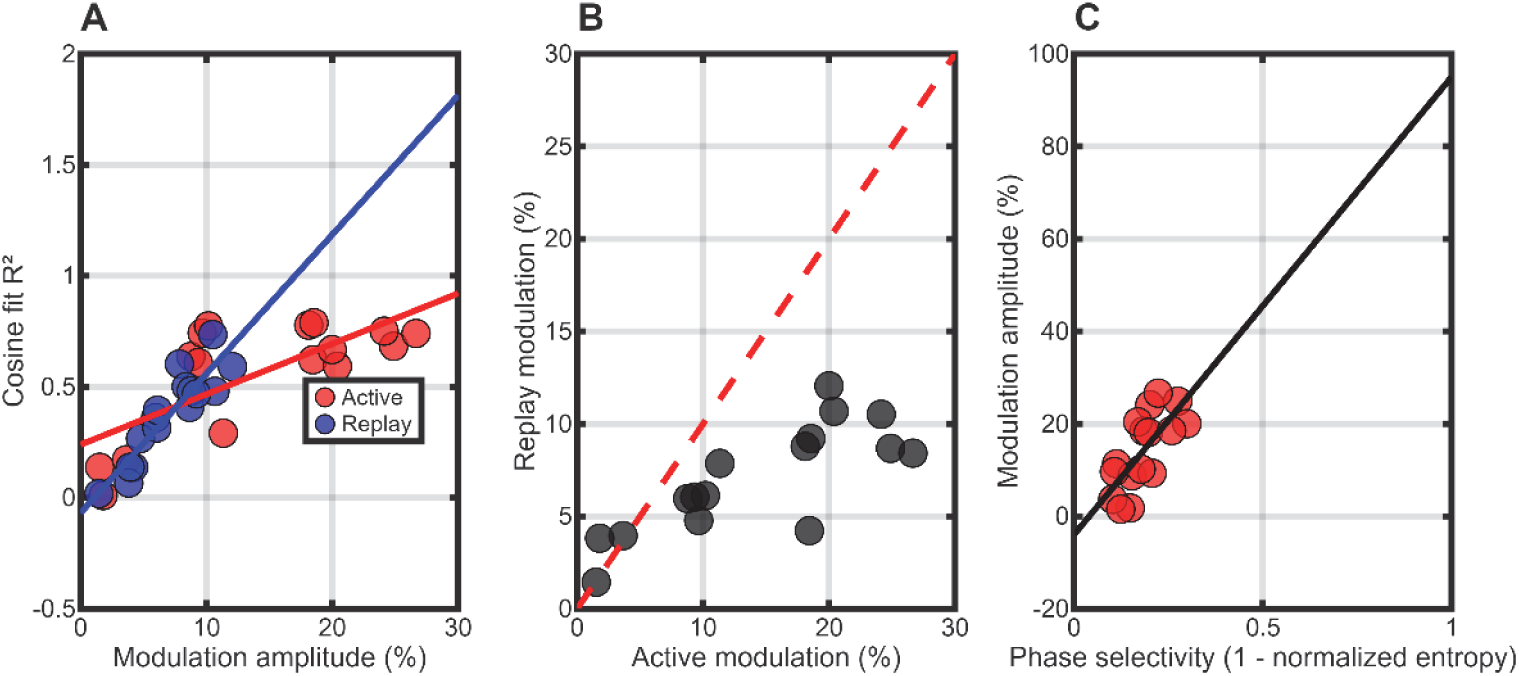
Relationships between modulation strength and tuning properties. **(A)** Correlation between modulation amplitude and cosine fit quality (R²) for active (red) and replay (blue) conditions. Both showed significant positive correlations (active: r = 0.73, p = 0.001; replay: r = 0.91, p < 0.001). **(B)** Individual consistency: comparison of modulation amplitudes during active and replay conditions. The red dashed line indicates identity. Most participants fell below the line, indicating stronger active modulation (r = 0.80, p < 0.001). **(C)** Phase selectivity (1 – normalized entropy) predicts modulation amplitude. Participants with more concentrated phase-specific modulation showed stronger effects (r = 0.72, p = 0.002).

### Modulation o alpha and beta

Figure 7 examines whether alpha and beta oscillations showed differential phase-dependent modulation across participants (n=16). Alpha oscillations exhibited bidirectional phase-dependent modulation (Figure 7A). Stimulation at phases 0°-135° produced a mean power increase of 21.0% (SEM = 19.9%), while phases 180°-315° produced a mean power decrease of - 13.3% (SEM = 10.8%; Figure 7A). Beta oscillations showed a different pattern (Figure 7B). Across all stimulation phases, beta power was predominantly suppressed, with a mean change of −9.3% (SEM = 9.6%; Figure 7B). No consistent phase-dependent boosting was observed for beta oscillations.

**Figure 7.**
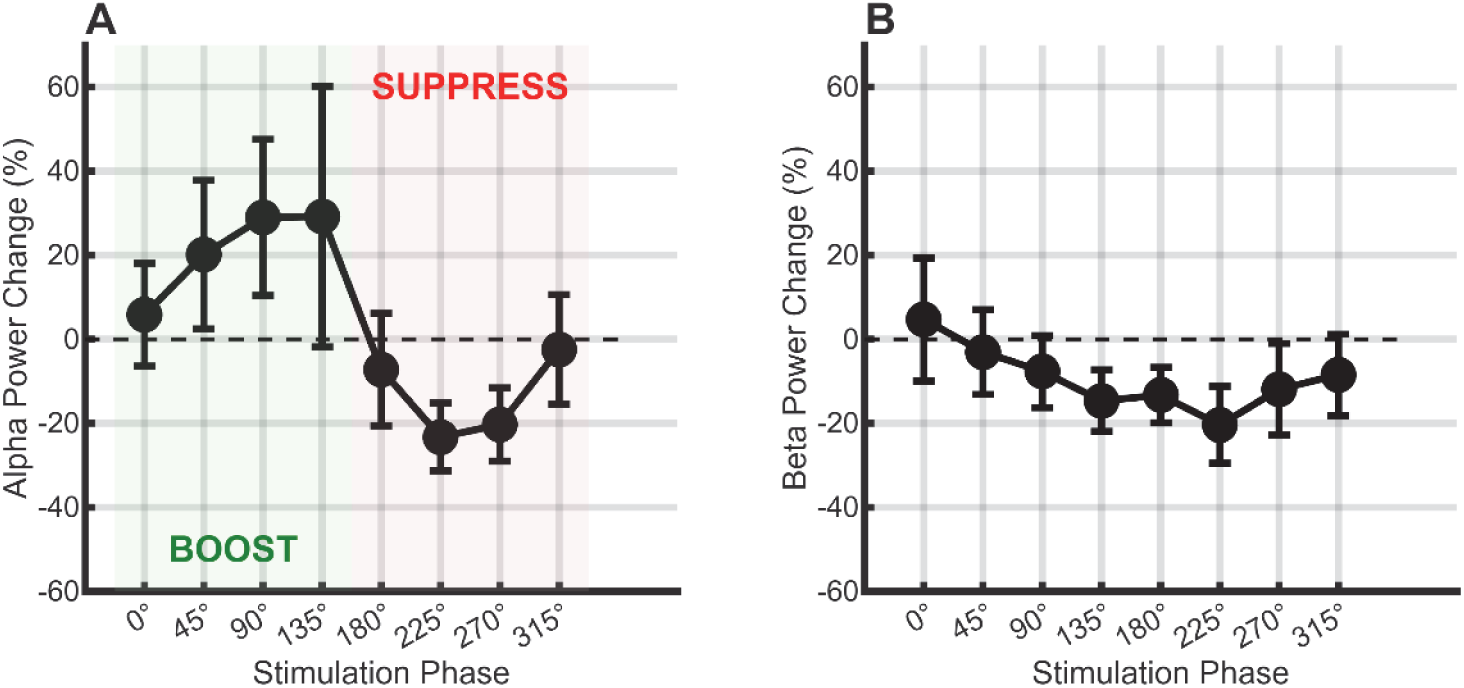
Alpha oscillations are bidirectionally modulated while beta oscillations are predominantly suppressed. **(A)** Alpha (8-12 Hz) power was bidirectionally modulated: phases 0°-135° increased power (mean: +21.0 ± 19.9%), while phases 180°-315° decreased power (mean: −13.3 ± 10.8%). **(B)** Beta (13-30 Hz) power showed unidirectional suppression across all phases (mean: −9.3 ± 9.6%), with no evidence of phase-dependent boosting.

### Dynamic requency tracking adapts to individual natural peaks

Figure 8 illustrates the critical importance of aligning stimulation with each individual’s natural peak frequency. A successful case of fixed-frequency targeting is shown, where the natural alpha peak at 10.1 Hz was closely matched by the target frequency (10.7 Hz; Figure 8A). Phase-dependent bidirectional modulation was observed, with the best boosting at 90° (+12.2%) and best suppression at 270° (−43.9%; modulation amplitude: 56.0%; Figure 8A). A failed case of fixed-frequency targeting is shown next. Here, the natural peak was in the beta range at 17.1 Hz, but stimulation targeted 8.5 Hz in the alpha range based on a priori assumptions (Figure 8B). At the target frequency, the best suppression was achieved at 315° (−19.9%), while the worst response (59.8% increase) occurred at 90°, resulting in a modulation amplitude of 79.7% (Figure 8B). Critically, different phase shifts produced power changes at different frequencies (phase-frequency coupling), and the natural beta peak remained largely unaffected (Figure 8B). Dynamic frequency tracking is shown in the final panel. The algorithm automatically identified the natural alpha peak at 10.1 Hz and targeted the full 8-30 Hz range (blue shaded area; Figure 8C). Clean bidirectional modulation was achieved without requiring a priori knowledge of the peak location, with best boosting at 45° (+18.0%) and best suppression at 225° (−54.7%; modulation amplitude: 72.7%; Figure 8C).

**Figure 8.**
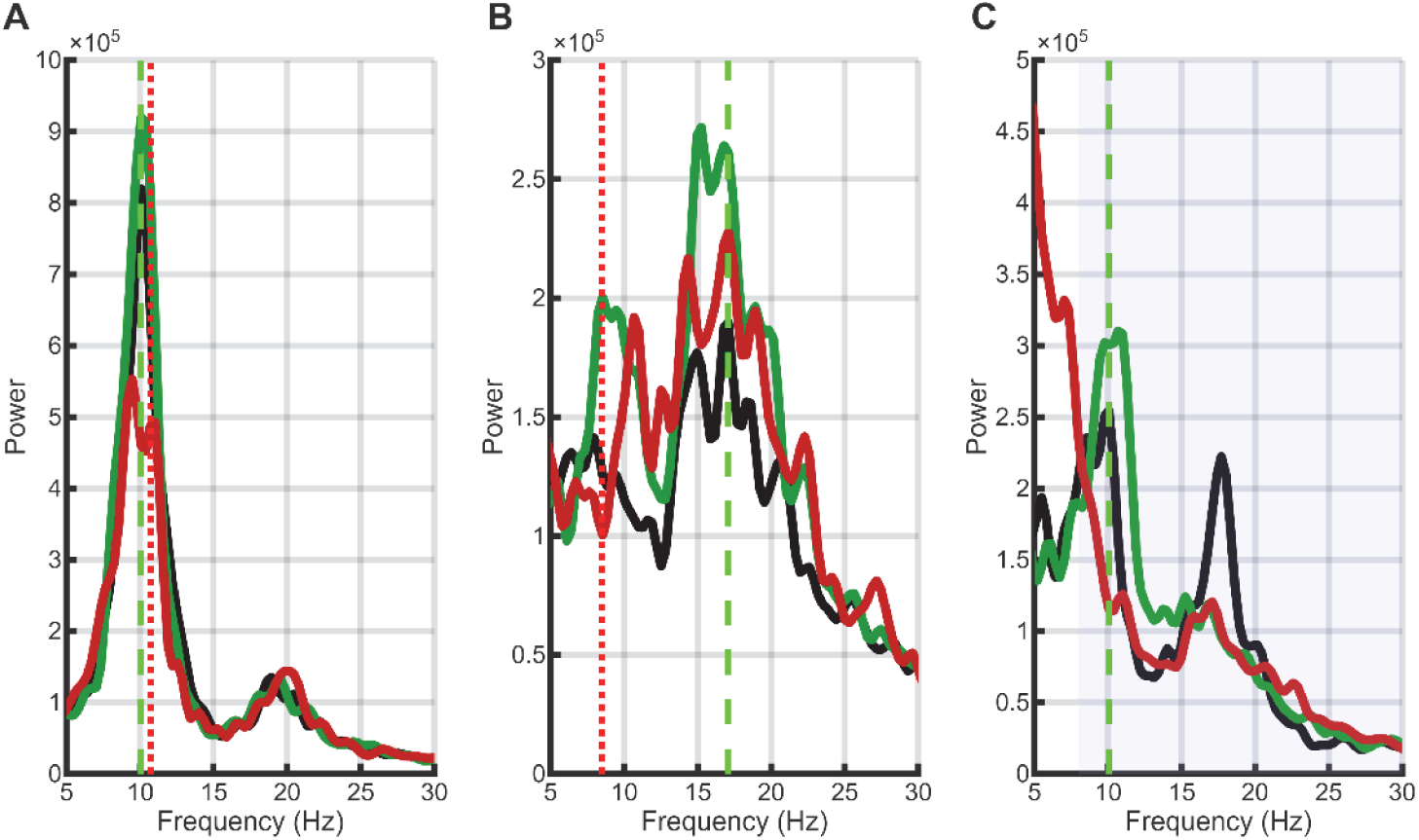
Fixed-frequency targeting requires alignment with natural peak; dynamic tracking automatically adapts. **(A)** Successful fixed-frequency targeting in one participant. The natural alpha peak at 10.1 Hz (green vertical line) was closely matched by the target frequency at 10.7 Hz (red vertical line). Phase-dependent bidirectional modulation was observed, with the best boosting at 90° (+12.2%, green) and best suppression at 270° (−43.9%, red; modulation amplitude: 56.0%). **(B)** Failed fixed-frequency targeting in another participant. The natural beta peak was at 17.1 Hz, but stimulation targeted 8.5 Hz (alpha range). At the target frequency, the best suppression was achieved at 315° (−19.9%), while the worst response (59.8% increase) occurred at 90°, resulting in a modulation amplitude of 79.7%. Note the scattered power changes across frequencies, indicating phase-frequency coupling when the target misses the natural peak. **(C)** Dynamic frequency tracking in a third participant. The algorithm automatically identified the natural alpha peak at 10.1 Hz and targeted the 8-30 Hz range (blue shaded area). Clean bidirectional modulation was achieved, with best boosting at 45° (+18.0%) and best suppression at 225° (−54.7%; modulation amplitude: 72.7%), without requiring a priori knowledge of the peak location.

## DISCUSSION

This study demonstrates that real-time, phase-locked peripheral nerve stimulation can selectively and causally modulate human sensorimotor oscillations. By combining a phase-modulated Poisson process with dynamic frequency tracking and statistically matched replay controls, we provide evidence that phase-dependent neuromodulation is not merely a by-product of stimulation statistics but reflects a genuine interaction between external input and the endogenous oscillatory state of the brain.

A central finding is that closed-loop phase-locked stimulation produces significantly stronger modulation than replay stimulation with identical temporal statistics (14.21% vs. 7.03%; p = 0.0003). This directly addresses a long-standing ambiguity in the field: whether apparent phase effects arise from true phase-dependent excitability or from residual correlations introduced by structured stimulation patterns (Duchet et al., 2023). By using replay controls derived from previously recorded neural activity, we ensured that stimulation rate, temporal distribution, and higher-order statistics were identical across conditions. The observed enhancement during active stimulation therefore isolates the causal contribution of real-time phase alignment. This provides strong evidence consistent with a causal role for the concept of phase-dependent excitability in human cortex, consistent with theoretical accounts in which oscillatory phase gates synaptic input efficacy (Jensen and Mazaheri, 2010; Kraus et al., 2016; Zanos et al., 2018).

A key methodological advance of this study is the use of a phase-modulated Poisson process. Traditional closed-loop stimulation paradigms often rely on periodic or quasi-periodic pulse trains, which risk entraining neural activity independently of endogenous dynamics (Fröhlich and McCormick, 2010; Reato et al., 2010). In contrast to fixed-interval stimulation, the stochastic timing used here reduces periodic entrainment while still delivering sufficient stimulation at the target phase to drive CNS responses. Moreover, single stimuli at a chosen phase may have too weak an effect; the modulated Poisson process provides greater power of input by concentrating multiple stimuli at the preferred phase, enabling robust probing of phase–excitability relationships.

The results reveal fundamental differences between alpha and beta oscillations in their phase– excitability profiles. Alpha activity exhibited clear bidirectional modulation, with phases 0°-135° producing a mean power increase of 21.0% and phases 180°-315° producing a mean decrease of 13.3%. This is consistent with models in which alpha oscillations implement rhythmic inhibition, creating alternating windows of high and low excitability (Jensen and Mazaheri, 2010). In contrast, beta oscillations showed predominantly unidirectional suppression across all phases (mean: −9.3%), suggesting a different underlying mechanism. Rather than reflecting symmetric excitability cycles, beta activity may represent a more stable or inhibitory network state, consistent with its proposed role in maintaining the current motor set (Engel and Fries, 2010), that is disrupted by peripheral input regardless of phase. This dissociation provides new insight into the functional roles of sensorimotor rhythms and indicates that phase-targeted stimulation may need to be tailored differently depending on the oscillatory regime being targeted.

Another important finding is the critical role of aligning stimulation with the individual’s natural peak frequency. Fixed-frequency targeting was effective only when the chosen frequency closely matched the endogenous oscillation. When this alignment failed, stimulation did not simply lose efficacy; instead, it produced complex phase-dependent effects distributed across frequencies—a phenomenon we describe as phase–frequency coupling. This suggests that phase is not an absolute quantity but is intrinsically linked to the oscillatory frequency at which it is defined. Misalignment between target and endogenous frequency therefore leads to ambiguous or distributed effects, rather than a simple reduction in modulation strength, consistent with the view that the effect of oscillatory phase is fundamentally frequency-specific (Fries, 2005).

The success of dynamic frequency tracking addresses this limitation directly. By continuously identifying the dominant frequency within 8-30 Hz on each processing cycle, the system ensured that phase estimation remained physiologically meaningful at all times. This approach eliminated the need for a priori band definitions and enabled robust bidirectional modulation across participants without prior knowledge of individual spectral profiles (mean modulation amplitude: 35.5%; active/replay ratio: 1.77). These findings argue strongly against the use of fixed canonical bands (e.g., alpha, beta) in closed-loop paradigms and instead support adaptive, data-driven targeting strategies (Haegens et al., 2014; van Ede et al., 2018).

Our closed-loop intervention elicited maximal oscillatory power enhancement at a mean phase of 70.2° (ascending phase approaching the peak of the sensorimotor oscillation), rather than at the trough (∼180°) often associated with maximal corticospinal excitability in transcranial magnetic stimulation (TMS) studies (Zrenner et al., 2018). This apparent discrepancy highlights that the relationship between oscillatory phase and neural responsiveness is not fixed, but depends on the measurement modality and the underlying neural processes being probed. MEG signals primarily reflect synaptic population activity, whereas TMS measures corticospinal output, and these signals are not necessarily phase-aligned.

Several factors may contribute to the observed phase offset. First, conduction delays between peripheral stimulation and cortical arrival (∼20 ms, as reflected by the N20m response) will introduce a systematic phase shift relative to the ongoing oscillation (Allison et al., 1989; Hari and Salmelin, 1997). Second, the phase at which afferent input most effectively interacts with ongoing activity may differ from the phase of maximal spontaneous excitability, particularly in distributed sensorimotor networks, consistent with models of phase-dependent gating of synaptic input (Jensen and Mazaheri, 2010). Third, median nerve stimulation at submaximal intensities engages both central sensory and motor pathways (Hamada et al., 2007), potentially biasing the effective interaction toward phases associated with sensorimotor integration rather than purely cortical excitability.

Taken together, these considerations suggest that the “optimal phase” for neuromodulation is not universal, but emerges from the interaction between stimulation modality, network dynamics, and signal type. This reinforces the importance of empirically determining effective phase relationships within each experimental and physiological context.

Our findings complement and extend previous work on brain state-dependent neuromodulation. Kraus et al. (2016) demonstrated that TMS applied during beta-band event-related desynchronization (ERD) elicited greater corticospinal excitation, whereas identical stimulation delivered outside this state elicited less excitation. Similarly, Zrenner et al. (Zrenner et al., 2018) showed that repetitive TMS timed to specific phases of the sensorimotor μ-rhythm induced phase-dependent plasticity effects. Our earlier work (Mitchell et al., 2007) further demonstrated that corticospinal responses to transcranial stimulation are modulated by ongoing oscillatory activity, particularly in the beta band, although this relationship is only partially predictive and depends on the signal modality. Together, these studies and the present results converge to the conclusion that the efficacy of stimulation is strongly dependent on the instantaneous brain state. However, they also highlight that the relationship between oscillatory phase and neural responsiveness is complex and not universally defined, varying across measurement modalities (EEG, EMG, MEG), stimulation approaches, and network dynamics (Bergmann et al., 2016). Our results extend this framework by demonstrating that real-time phase-targeted peripheral stimulation can causally modulate oscillatory activity when aligned with the endogenous rhythm, and by providing a methodological framework that isolates true phase-dependent effects from stimulation statistics.

At a broader level, the present results establish several principled guidelines for closed-loop neuromodulation. First, stimulation timing should avoid regular periodicity to prevent artificial entrainment; stochastic Poisson stimulation provides a physiologically valid alternative. Second, phase estimation must be tied to the current dominant frequency rather than predefined canonical bands. Third, causal inference requires controls that match stimulation statistics while removing real-time contingency; replay controls provide a rigorous solution. Finally, the functional effects of stimulation depend on the specific oscillatory regime (alpha vs. beta), implying that protocols must be tailored to the underlying neural dynamics.

Several limitations should be considered. The behavioural task was relatively simple and designed primarily to stabilize oscillatory activity; future work should investigate how phase-targeted stimulation influences behaviour and motor performance. In addition, while peripheral nerve stimulation provides a safe and well-controlled input pathway, it engages multiple downstream pathways, making it difficult to isolate the precise neural mechanisms underlying the observed effects. The precision of phase estimation is inherently limited by filtering, buffering, and sensor-level signal mixing, although any residual imprecision would be expected to reduce, rather than inflate, phase specificity.

Future studies could extend this framework in several directions. Combining phase-targeted stimulation with behavioural paradigms may reveal whether modulating oscillatory phase can enhance motor learning or cognitive performance. Integration with other stimulation modalities, such as transcranial magnetic or electrical stimulation, could test whether the same principles generalize across input pathways. Finally, applying this approach in clinical populations may offer new strategies for disorders characterized by abnormal oscillatory dynamics, such as Parkinson’s disease (Hammond et al., 2007), essential tremor, or post-stroke motor deficits.

In conclusion, this study provides direct evidence that phase-targeted, stochastic closed-loop stimulation can selectively modulate human brain oscillations in a causal and frequency-specific manner. By demonstrating the importance of phase alignment, frequency matching, and stimulation statistics, we establish a unified and physiologically grounded framework for closed-loop neuromodulation. These findings not only clarify the mechanisms of phase-dependent stimulation but also lay the foundation for personalized, adaptive neurostimulation strategies in both research and clinical applications.

## Author Contributions

B.Z. designed the research, performed the experiments, analysed the data, wrote the first draft of the paper, and edited the paper. S.N.B. designed the research, performed the experiments, contributed analytic tools, interpreted the data, and edited the paper.

## Acknowledgement

The authors thank Dr Sian Worthen for her invaluable support with MEG and her continuous technical assistance throughout the study. This work was supported by an internal research grant from Aston University.

